# A Bacterial Expression Vector Archive(BEVA) for flexible modular assembly of Golden Gate-compatible vectors

**DOI:** 10.1101/392423

**Authors:** Barney A. Geddes, Marcela A. Mendoza-Suárez, Philip S. Poole

## Abstract

We present a Bacterial Expression Vector Archive (BEVA) for the modular assembly of bacterial vectors compatible with both traditional and Golden Gate cloning, utilizing the Type IIS restriction enzyme Esp3I. Ideal for synthetic biology and other applications, this modular system allows a rapid, low-cost assembly of new vectors tailored to specific tasks. To demonstrate the potential of the system three example vectors were constructed and tested. Golden Gate level 1 vectors; pOGG024, with a broad-host range and high copy number was used for gene expression in laboratory-cultured *Rhizobium leguminosarum*, and pOGG026, with a broad-host range a lower copy number and excellent stability, even in the absence of antibiotic selection. The application of pOGG026 is demonstrated in environmental samples by bacterial gene expression in nitrogen-fixing nodules on pea plants roots formed by *R. leguminosarum*. Finally, the level 2 cloning vector pOGG216 is a broad-host range, medium copy number, for which we demonstrate an application by constructing a dual reporter plasmid expressing green and red fluorescent proteins.

**IMPORTANCE:** Modular assembly is powerful as it allows easy combining of different components from a library of parts. In designing a modular vector assembly system, the key constituent parts (and modules) are; an origin of plasmid replication, antibiotic resistance marker(s), cloning site(s), together with additional accessory modules as required. In an ideal vector, the size of each module would be minimized, and this we have addressed. We have designed such a vector assembly system by utilizing the Type IIS restriction enzyme Esp3I and have demonstrated its use for Golden Gate cloning in *Escherichia coli*. An important attribute of this modular vector assembly is that using the principles outlined here, new modules for specific applications, e.g. origin of replication for plasmids in other bacteria, can easily be designed. It is hoped that this vector construction system will be expanded by the scientific community over time by creation of novel modules through an open source approach.

---

Over the last decade, synthetic biology has emerged as a powerful tool for biotechnology and basic science research (1). Coupled with rapidly improving DNA synthesis technologies, one of the key advances that has potentiated the rapid expansion of synthetic biology is new DNA assembly technologies (2). The ability for synthetic biologists to rapidly assemble and test new designs at low cost is invaluable to progress in a wide-range of life sciences research. Applications for this range from investigating new antibody production with higher success rates, to increased food production minimizing the carbon footprint with the effective transfer of nitrogen fixation between bacterial species (3).

Golden Gate cloning is a DNA assembly technology that utilizes type IIS restriction enzymes to assemble multiple fragments of DNA in a linear order (4). Since type IIS restriction endonucleases cleave outside of their recognition site, the nucleotides in the cut-site of the enzyme are not discriminated and can be tailored to suit. This permits assembly of multiple fragments of DNA in a directional, linear order by using unique cut/ligation sites between each fragment, all of which can be cut by the same type IIS restriction enzyme. The two most commonly used Golden Gate enzymes are BsaI and BpiI, each has a 6 base-pair (bp) recognition site and a 4 bp sticky-end cut site (4). The ability of these enzymes to cleave outside of their recognition site and careful design of compatible overhanging sequences allows repeated rounds of cutting followed by ligation to proceed towards a stable assembly product which remains undigested as it lacks enzyme recognition sites (due to the inward orientation of the type IIS recognition sites at the far 5’ and 3’ ends of modules) (5). This approach has given a massive improvement in the efficiency of assembly compared to traditional rounds of restriction/ligation cloning and has been used to assemble at least nine fragments in linear order, with 90% of transformed bacterial colonies containing correct products (5).

Golden Gate cloning platforms have been developed for plants, fungi and the bacterium *Escherichia coli* (6–9). However, in a wide-range of bacteria, vectors for cloning using Golden Gate tools are not available. The ability to use the most up-to-date molecular tools is crucially important for agricultural biotechnology. Here we describe the development of a system for modular assembly of Golden Gate-compatible cloning broad-host range vectors from a library of vector parts. As these vectors are able to replicate in *E. coli*, this facilitates easy genetic manipulation since following plasmid construction, as the plasmids constructed can be transferred easily to other bacterial species. We demonstrate the efficacy of this system by constructing several vectors, analogous to useful traditional cloning broad-host range vectors but that are now Golden Gate-compatible: level 1 cloning broad-host range vectors pOGG024, a medium copy plasmid for gene expression for *in vitro* grown bacteria, and pOGG026, a low copy number stable vectors for stable environmental gene expression in environmental samples where no antibiotic selection is applied. We also describe the construction through this modular vector system of a level 2 broad-host range vector pOGG216.

## RESULTS AND DISCUSSION

### Modular vector assembly

A new standardized system for vector assembly was designed based on Golden Gate cloning and the MoClo system described by Weber *et al.* 2011 (4). We have designed a bacterial vector assembly system with a series of discrete key modules. In order to do this we took advantage of the modularization and miniaturization of bacterial vector modules in the Standard European Vector Archive (SEVA) (10, 11). In SEVA these modules are arranged in a predefined order and assembled using standard restriction enzymes and standard restriction/ligation reactions. We have conserved this organization in the modular vector assembly system: position 1 is the cloning site(s), position 2 is the antibiotic resistance cassette, position 3 is origins of replication and transfer.

Three more positions (4 to 6) have been reserved for additional accessory modules for specific applications. Endlinkers containing terminators (ELT) are used to circularize the plasmid by connecting the final position used to position 1 (Fig. 1A). We incorporated Golden Gate level 1 and level 2 cloning sites into position 1 to make them compatible with the MoClo Golden Gate cloning system (Fig. 1B). Each vector construction module is flanked with inward-facing recognition sites for the type IIS restriction enzyme Esp3I and the cut sites, giving overhangs with sequences specific for each module (fusion sites shown in Table 1), allow directional linear assembly in a one-pot reaction such as used in Golden Gate cloning (4).

**FIG 1.**
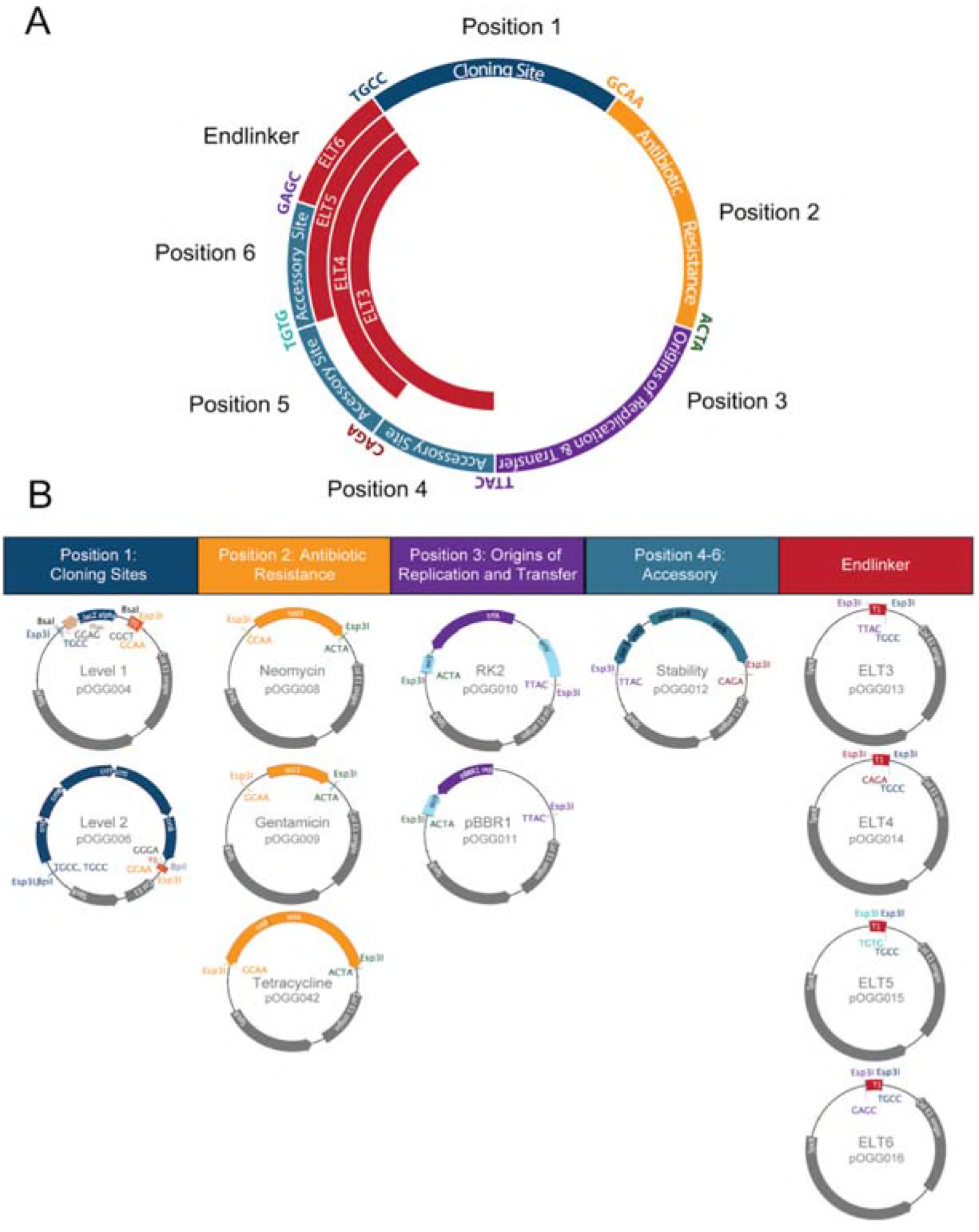
Bacterial Expression Vector Archive (BEVA) system for the modular assembly of bacterial vectors. (A) Vector modules are designated a ‘position’ and combined in the vector construction as shown. Sequences (5’ → 3’) used to anneal the fragments in order are shown (colour-coded) at junctions between modules and remain in the final construct as a scar. (B) Modules at each position. Position 1: Cloning Sites-Golden Gate level 1 and Golden Gate level 2; position 2: Antibiotic resistance-neomycin-, gentamicin- and tetracycline-resistance cassettes; position 3: Origins of replication and transfer – RK2 with *oriT* and pBBR1 with *oriT;* positions 4-6: Accessory modules – *par* locus for stable plasmid maintenance in the absence of antibiotic selection. Endlinker modules were designed to circularize the plasmid.

**Table 1.** Design of modules for vector construction.

| Module | Description of module | Module size (kb) | Cloned module | 5'-side (5' → 3') | 3'-side (5' → 3') |
| --- | --- | --- | --- | --- | --- |
| **Position 1: Cloning sites** |  |  |  | TGCC | GCAA |
| pLVC-P1-Lv1 | Golden Gate Level 1 cloning site with cloned *lacZ* | 2601 | pOGG004 |  |  |
|  | Golden Gate Level 1 cloning site | 2420 | pOGG005 |  |  |
| pLVC-P1-Lv2 | Golden Gate Level 2 cloning site | 5729 | pOGG006 |  |  |
| **Position 2: Antibiotic resistance** |  |  |  | GCAA | ACTA |
| pLVC-P2-neo | Neomycin/kanamycin-resistance | 936 | pOGG008 |  |  |
| pLVC-P2-gent | Gentamicin-resistance | 813 | pOGG009 |  |  |
| pLVC-P2-tet | Tetracycline-resistance | 1964 | pOGG042 |  |  |
| **Position 3: Origins of replication and transfer** |  |  |  | ACTA | TTAC |
| pLVC-P3-RK2 | RK2 (IncP) origin of replication for low copy number plasmid, *oriT* from *E. coli* for plasmid transfer | 2485 | pOGG010 |  |  |
| pLVC-P3-pBBR1 | pBBR1 origin of replication for medium copy number plasmid, *oriT* from *E. coli* for plasmid transfer | 1784 | pOGG011 |  |  |
| **Position 4: Accessory** |  |  |  | TTAC | CAGA |
| pLVC-P4-par | Partition genes (*parABCDE*) for plasmid stability in absence of antibiotic selection | 2408 | pOGG012 |  |  |
| **Position 5: Accessory** |  |  |  | CAGA | TGTG |
| **Position 6: Accessory** |  |  |  | TGTG | GAGC |
| **Endlinker** |  |  |  |  |  |
| pLVC-ELT-3 | Endlinker to circularize plasmid after position 3 | 113 | pOGG013 | TTAC | TGCC |
| pLVC-ELT-4 | Endlinker to circularize plasmid after position 4 | 113 | pOGG014 | CAGA | TGCC |
| pLVC-ELT-5 | Endlinker to circularize plasmid after position 5 | 113 | pOGG015 | TGTG | TGCC |
| pLVC-ELT-6 | Endlinker to circularize plasmid after position 6 | 113 | pOGG016 | GAGC | TGCC |

### Vector module design

We assigned modules a universal position for linear assembly based on function; position 1 – cloning site(s), position 2 – antibiotic resistance, position 3 – origins of replication and transfer, with positions 4–6 reserved for accessory modules (Fig 1, Table 1). Modules at each position (1 to 6 and Endlinkers, Table 1) have inward-facing Esp3I sites which on digestion give 4 bp-overhangs. These overhangs are able to anneal with those from neighbouring modules in a linear fashion and, on vector assembly, give 4 bp motifs which remain at the join between modules (all motifs for each junction are listed in Table 1, Fig. 1). An advantage of modular assembly is the ability to utilize minimal parts which exclude any non-functional DNA. This allows construction of vectors with the same key features as those currently widely adopted, but the resulting plasmid is significantly smaller in size. The reduced size enhances the ease of DNA manipulation and allows accommodation of larger cloned inserts. To this end, several modules (positions 2 and 3) used the minimal units of antibiotic-resistance modules and origins of replication/transfer previously defined by SEVA (10, 11) (Table 1).

Most parts were constructed by DNA synthesis by Invitrogen and supplied in the GeneArt cloning vector pMS (Invitrogen, Thermo Fisher Scientific Inc.). As an alternative some vector modules were constructed by PCR amplification from commonly used vectors (see Table 3). In several cases, this was a cheaper and faster option than the synthesis of the DNA fragments. Modules synthesized by Invitrogen were domesticated prior to DNA synthesis by removal of BsaI, BpiI, DraIII and Esp3I (Golden Gate cloning restriction enzyme sites) by altering a single nucleotide in the recognition site. In an open reading frame (ORF) a neutral change was made in the wobble base of a codon, such that the same amino acid was encoded. Outside of an ORF, a transition mutation was introduced.

The position 1 modules we designed are based on level 1 and level 2 cloning sites pL1F-1 and pL2V-1 from the MoClo system of Weber *et al.* (4) (Table 1). The level 1 cloning site in pOGG004 contains a *lacZα* fragment that is replaced following Golden Gate cloning following the MoClo system, resulting in blue to white colony colour selection when plated on appropriate media. Since *lacZα* still contains a polylinker, vectors constructed with the level 1 Golden Gate cloning site are also flexiblefor use in traditional cloning utilizing unique restriction sites in the polylinker and blue-white selection. The level 2 golden gate cloning site encodes a canthaxanthin biosynthesis operon that is replaced upon successful level 2 cloning resulting in a transition from an orange colony colour to white or blue. Both position 1 modules include a *rho*-independent T_0_ lambda phage transcription terminator (12) included at their 3’-end.

We note that since the level 1 Golden Gate cloning site part (pOGG004) is maintained in a high copy number vector backbone, it can be cloned into directly using BsaI as described here and utilized as a high copy plasmid for recombinant protein expression in *E. coli*. We synthesised (pOGG005), with an Ampicillin resistant backbone for this purpose, to be compatible with traditional systems for recombinant protein production.

For position 2 (antibiotic resistance), gentamicin- and neomycin-resistance modules were based on the minimal resistance cassettes from SEVA(11). The 5’-end SwaI and 3’-end PshAI sites that flanked the SEVA modules were replaced with inward-facing Esp3I sites. We found that the SEVA constitutive *tetA* gene conferred insufficient levels of tetracycline resistance to our organisms of interest (data not shown), therefore we designed a more robust tetracycline-resistance module based on the *tetAR* resistance module from pJP2 (13). To preserve a minimal size, only DNA between the stop codons of the divergently transcribed *tetA* and *tetR* was used and their orientation conserved (Fig. 1B). In position 3 (origins of replication and transfer), modules containing pBBR1 and RK2 origins of replication were designed based on SEVA modules (11), but with inward facing Esp3I sites replacing PshAI and AscI sites. An origin of transfer (*oriT*), to permit conjugation from *E. coli* into target strains, was included at the 5’-end. At position 4, an accessory module for stable plasmid maintenance in the absence of antibiotic selection was designed. Based on the *par* locus from pJP2, which has previously been shown to render stable plasmid maintenance in the absence of antibiotic selection (13), *parABCDE*, as well as 107 bp downstream of *parE* (at 5’-end) and 139 bp downstream of *parA* (at 3’-end) (Fig 1B) was oriented such that the *parE* was transcribed towards position 3, while *parA* was at the 3’-end, transcribed towards position 1 (Fig. 1B).

Endlinker (ELT) modules were designed to complete assembly by circularizing the plasmid, either by linking position 3 (origin of replication), or accessory modules at positions 4, 5 or 6 to position 1 (cloning sites). Endlinker modules contain the *E. coli* T_1_ *rrnBT1* terminator to help buffer any transcription effects from the cloning site (12).

### Design and construction of level 0 open reading frame parts

Golden Gate cloning is often used in a hierarchical assembly where level 0 parts (genetic features controlling a gene’s transcription, translation and the termination of transcription) are assembled in a single one-pot reaction (referred to as level 1 assembly) (4). In order to test the vectors that we constructed, we adapted several parts that are routinely used in our lab, including promoters with ribosome binding sites (PU modules), ORFs (SC modules) and a terminator (T module). These parts were synthesized as described for vector construction modules, except that they were assembled with inward-facing BsaI sites that cut appropriate sequences for level 1 assembly (Table 3).

We adapted several promoters routinely used for gene expression in rhizobia or *E. coli* for use as PU modules (Table 3). These are the constitutive neomycin cassette promoter (pNptII) (14), T7RNAP promoter repressible by LacI (p*T7lacO*,)(15), the Lac promoter from *E. coli* (*pLac*) (16), the taurine-inducible promoter of *S. meliloti* (*pTau*) (17) and the symbiosis-induced *nifH* promoter from *Rhizobium leguminosarum* biovar *vicia* 3841 (Rlv3841pNifH) (18). For all promoters, the nucleotide immediately upstream of the ATG was domesticated to an A to construct the 5’-AATG-3’ cut-site such that the ATG encodes the start codon of the ORF to be expressed.

For SC modules (Table 3), we utilized several different reporter genes to demonstrate vector function. These include genes encoding superfolder green fluorescent protein (sfGFP) (19), *E. coli* β-glucoronidase (*gusA*) and *Pyrococcus furiosus* thermostable β-glucosidase/ β-galactosidase (*celB*) (20, 21).

Domestication and gene synthesis were used to produce SC modules of sfGFP and *celB*, and *gusA*. All SC modules were constructed flanked by inward facing BsaI restriction site cutting 5’-AATG-3’ at the 5’ end and GCTT at the 3’ end. In all cases the final three nucleotides of the AATG scar encode the start codon for the ORF and the GCTT scar immediately follows the stop codon.

For use as a transcriptional terminator, the *rrnBT1* transcriptional terminator with flanking regions from pLMB509 (17) (9 bp upstream and 84 bp downstream) were synthesised and flanked by BsaI sites cutting GCTT and CGCT on the 5’ and 3’ of the open reading frame. All the novel BEVA parts, vectors and plasmids are available through Addgene (http://www.addgene.org/ Addgene ID is shown in Table 2).

**Table 2.**
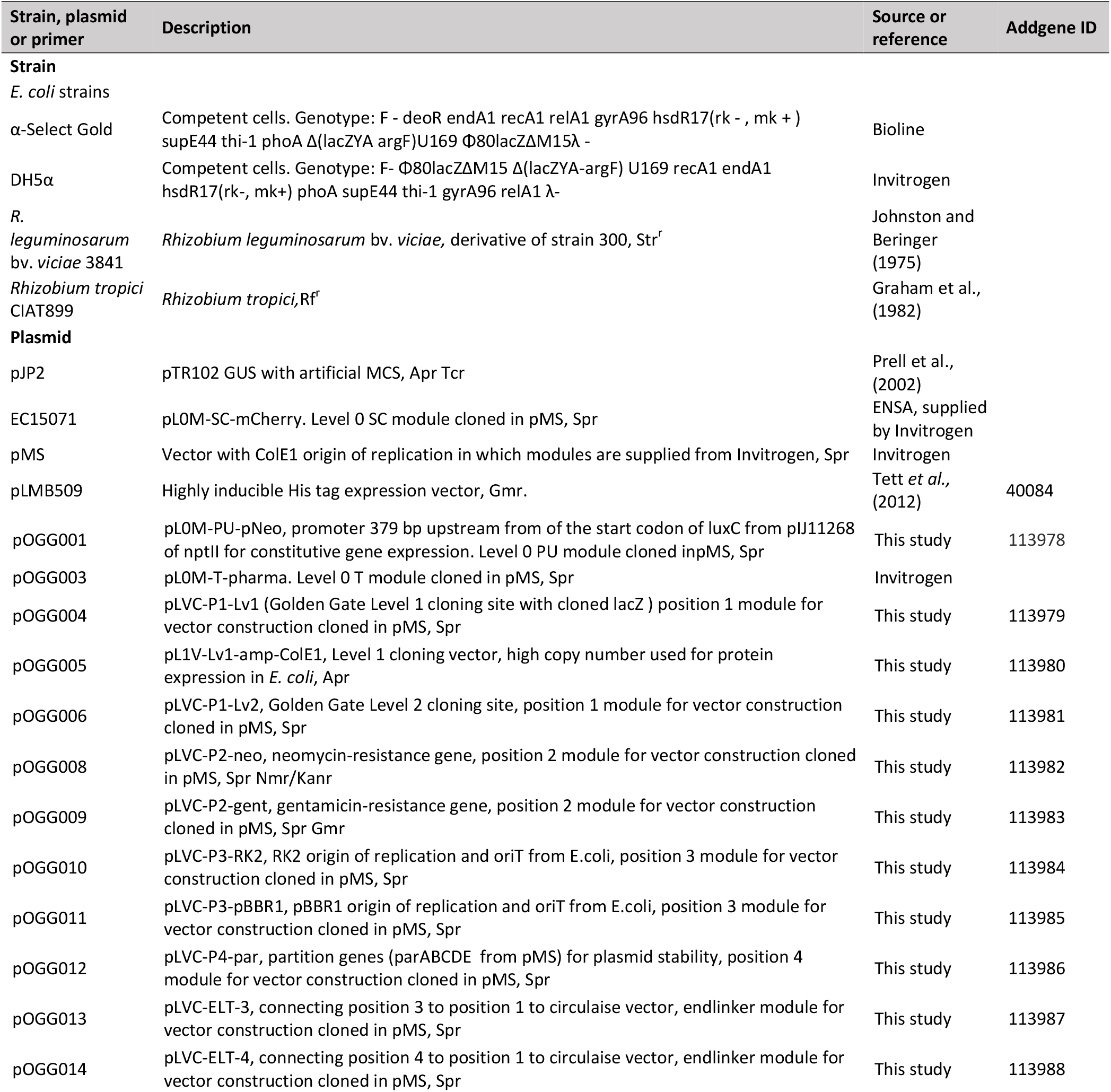
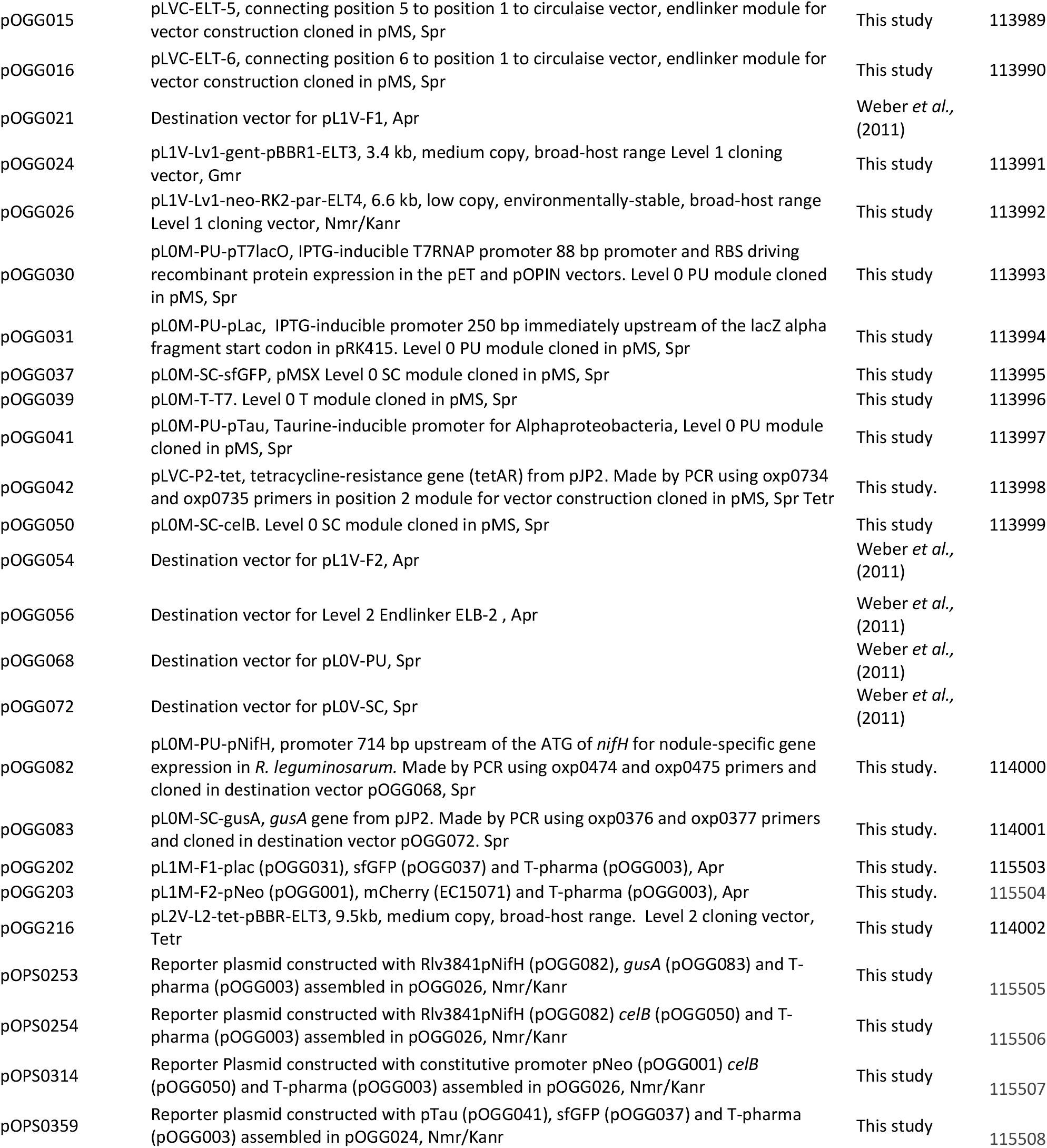
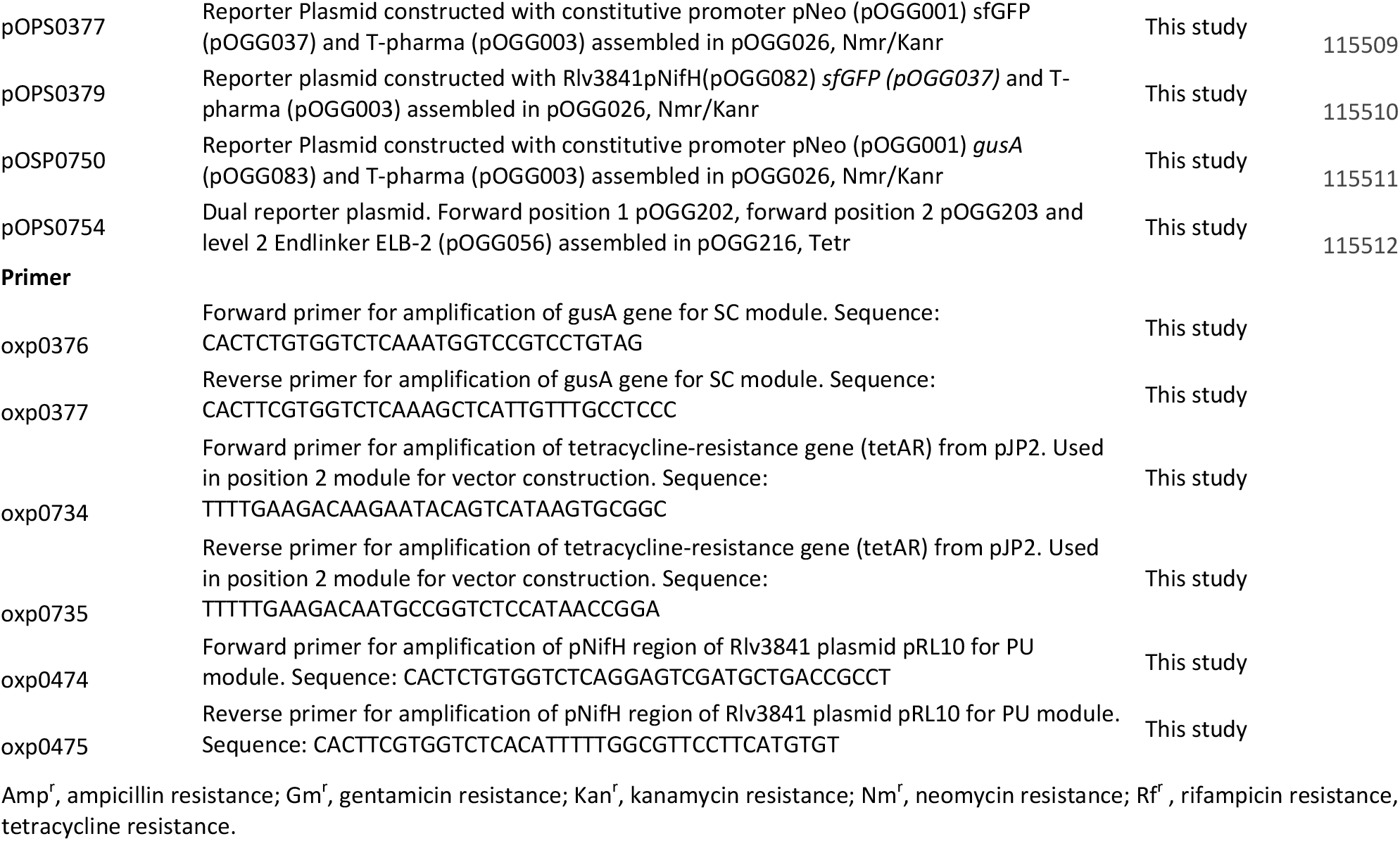
Strains, plasmids and primers used in this study. Amp^r^, ampicillin resistance; Gm^r^, gentamicin resistance; Kan^r^, kanamycin resistance; Nm^r^, neomycin resistance; Rf^r^, rifampicin resistance, Tet^r^, tetracycline resistance.

### Vectors for Golden Gate level 1 cloning in bacteria

Vector assembly reactions were performed by selecting appropriate modules in each position (Fig. 1, summary of the sequences used to combine the modules which remain at their junctions and fusion sites are given in Table 1).

To demonstrate the flexibility of the system, two vectors for level 1 cloning, analogous to those used for traditional cloning and DNA manipulation in Gram-negative bacteria, were constructed. Plasmid pOGG024 (Fig. 2A, Table 2), a 3.4 kb level 1-compatible vector for bacterial gene expression, was made by adding the following plasmids (listed in Table 3) to a one-pot reaction (digestion with Esp3I and ligation) to combine the following modules: position 1; level 1 cloning site from pOGG004, position 2; gentamicin resistance cassette from pOGG009, position 3; pBBR1-based origin of replication from pOGG011 and the appropriate Endlinker (ELT3) from pOGG013 (Table 1).

**FIG 2.**
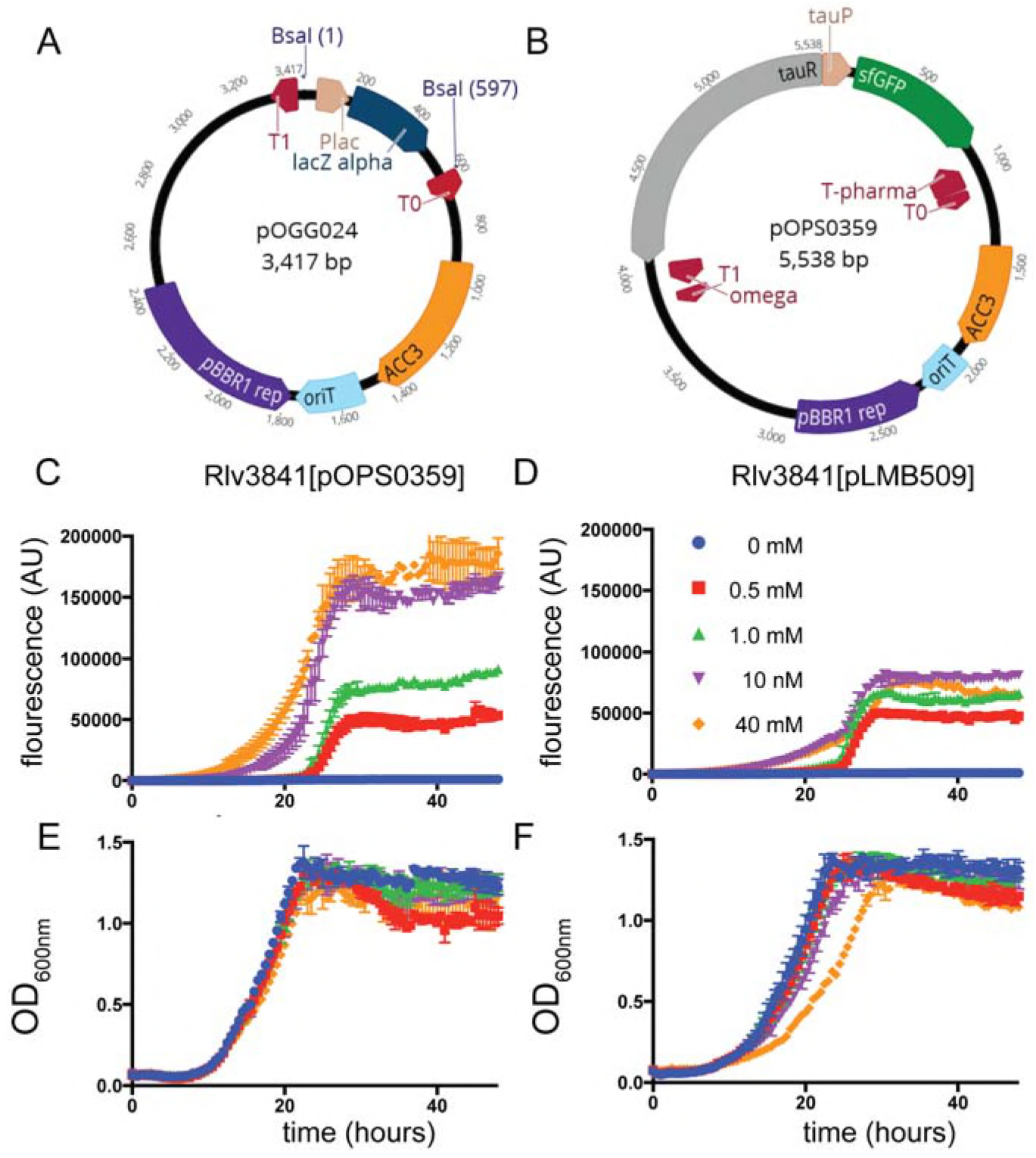
A) Schematic map of pOGG024 level 1 broad-host range, high copy number vector constructed with BEVA for free-living gene expression in laboratory. B) Schematic map of plasmid pOPS0359 containing a cloned gene sf-GFP under the control of a taurine-inducible promoter from *Sinorhizobium meliloti* into pOGG024. C-F) Evaluation of pOPS0359 performance as robust free-living gene expression compared to the functionally analogous pLMB509 plasmid. C-D) Florescence detection from GFP expression and E-F) Fitness performance by measurement of OD_600_ when grow without (0 mM) or with (0.5, 1.0, 10 or mM) taurine.

Plasmid pOGG026 (6.6 kb, Fig. 3A, Table 2) was constructed by combining the following modules (Table 3) in a one-pot reaction (digestion with Esp3I and ligation): position 1; level 1 cloning site from pOGG004, position 2; neomycin resistance gene from pOGG008, position 3; RK2-based origin of replication from pOGG010, position 4; *parABCDE* genes which confer stability in the absence of antibiotics from pOGG012 and the appropriate Endlinker (ELT4) from pOGG014 (Table 1).

**FIG 3.**
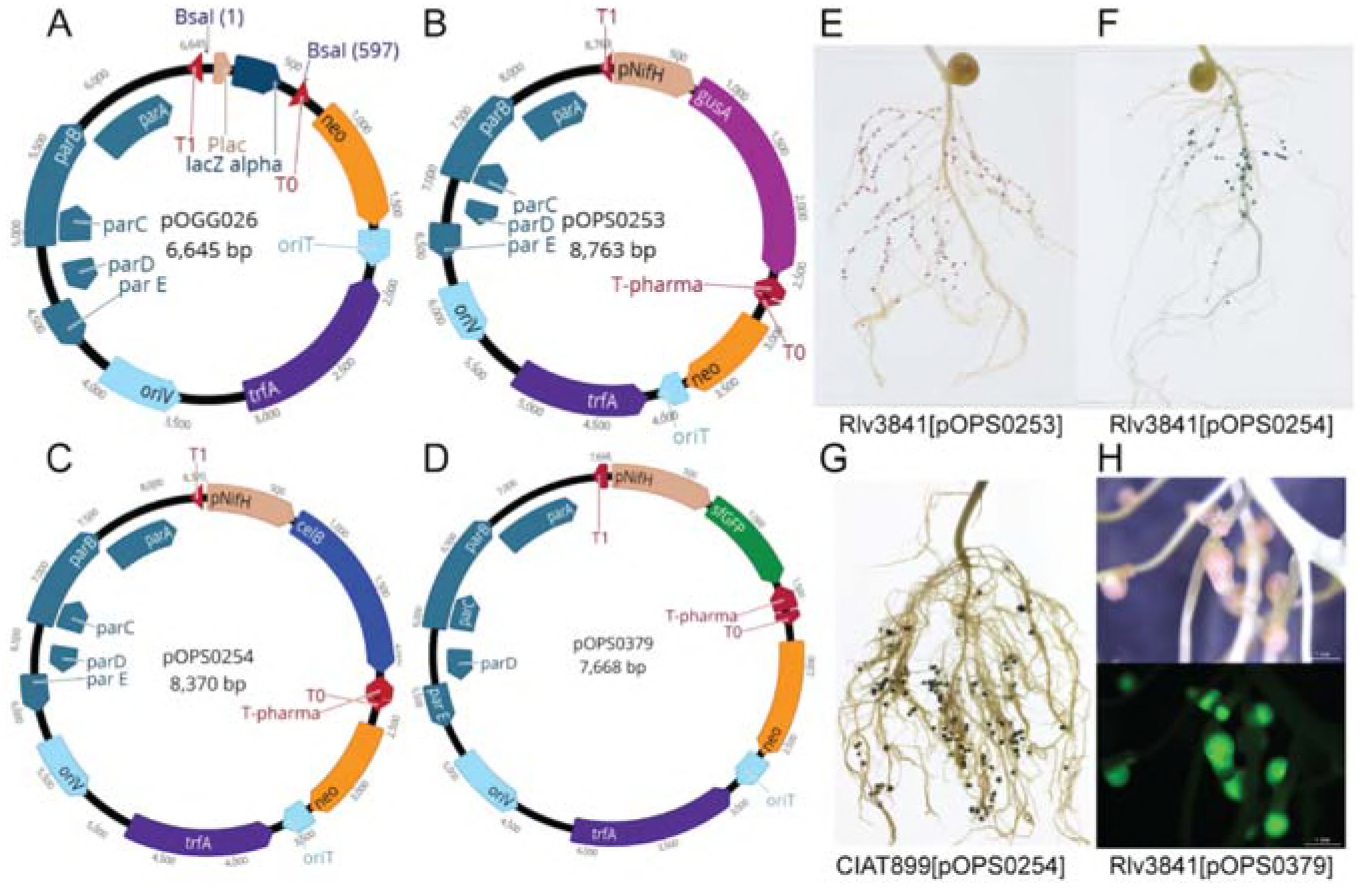
A) Schematic map of pOGG026 level 1 broad-host range, lower copy number vector constructed with BEVA for stable environmental gene expression in the absence of antibiotic selection. B-D) Schematic map of plasmids containing a cloned B) *gusA* C) *celB* and D) sfGFP gene under the control of *nifH* gene promoter (pNifH). E) Pea nodules formed by Rlv3841[pOPS253] stained with Magenta-glc. Marker gene *gusA* is just expressed in nodules. F) Pea nodules formed by Rlv3841[pOPS254] and G) bean nodules formed by CIAT899[pOPS253] stained with X-gal after thermal treatment. Marker gene *celB* is just expressed in nodules. H) Pea nodules formed by Rlv384[pOPS0379], bright-field (top) and fluorescence (GFP filter) (bottom) images. GFP expression is exclusively found in the nitrogen-fixing zone.

**Table 3.** Level 0 modules for cloning in a Level 1 vector. All modules were supplied cloned in Sp^r^ plasmid pMS except for pOGG082 and pOGG083. Following BsaI digestion of cloned modules and a Level 1 vector, the resulting overhangs are used to anneal the fragments in a linear fashion and the sequences shown remain between modules upon ligation.

| Module | Description of module | Module size (bp) | Cloned module | 5'-side (5' → 3') | 3'-side (5' → 3') |
| --- | --- | --- | --- | --- | --- |
| **PU modules: promoters with ribosome-binding sites** |  |  |  | GGAG | AATG |
| pL0M-PU-pNeo | Promoter upstream of *nptII* for constitutive gene expression | 386 | pOGG001 |  |  |
| pL0M-PU-pT7lacO | IPTG-inducible T7RNAP promoter | 95 | pOGG030 |  |  |
| pL0M-PU-pLac | IPTG-inducible promoter | 258 | pOGG031 |  |  |
| pL0M-PU-pTau | Taurine-inducible promoter for Alphaproteobacteria | 1811 | pOGG041 |  |  |
| pL0M-PU-pNifH | Promoter upstream of *nifH* for nodule-specific gene expression in *R. leguminosarum* | 721 | pOGG082 |  |  |
| **SC modules: ORFs** |  |  |  | AATG | GCTT |
| pL0M-SC-mCherry | Red fluorescence protein mCherry, reporter of gene expression | 716 | EC15071 |  |  |
| pL0M-SC-sfGFP | Superfolder GFP, stable and bright reporter of gene expression | 722 | pOGG037 |  |  |
| pL0M-SC-celB | Thermostable beta-galactosidase, reporter of gene expression | 1424 | pOGG050 |  |  |
| pL0M-SC-gusA | Beta-glucosidase gene, reporter of gene expression | 1817 | pOGG083 |  |  |
| **T modules: terminators** |  |  |  | GCTT | CGCT |
| pL0M-T-pharma | Pharmacia terminator (Invitrogen) | 188 | pOGG003 |  |  |
| pL0M-T-T7 | Terminator for T7RNAP | 55 | pOGG039 |  |  |

### Gene expression using vector pOGG024

In previous work we described the construction of pLMB509 (17), a plasmid which has a taurine-inducible promoter upstream of a gene encoding GFP (*gfpmut3.1*).To evaluate the performance of vector pOGG024 through comparison with the fully-characterized pLMB509, we assembled plasmid pOPS0359 by cloning a taurine-inducible promoter upstream of superfolder GFP (sfGFP) to create a functionally analogous plasmid i.e. taurine-inducible expression of GFP (Fig. 2B). To do this, a Golden Gate level 1 cloning reaction was performed with components (listed in Table 3) in a one-pot reaction (digestion with BsaI and ligation); vector; pOGG024, PU module; *Sinorhizobium meliloti* taurine promoter from pOGG041, SC module; sfGFP from pOGG037, T module; rrnBT1 terminator from pOGG003 to create pOPS0359. Taurine-dependent GFP expression of plasmids pOPS0359 and pLMB509 in a Rlv3841 background is shown in Fig. 2C-D. Under conditions of 0 to 40 mM taurine they showed similar growth (Fig 2E-F) and gave fluorescence profiles indicating induction of GFP throughout the exponential phase. Although similar GFP induction profiles were seen, pOPS0359 showed significantly higher levels of overall fluorescence, especially at 10 and 40 mM taurine. As the plasmids share the same origin of replication (pBBR1) there are two factors that are likely to have had an influence on this; the brighter fluorescence of sfGFP (19) (pOPS0359) compared to that of GFPmut3.1 (pLMB509), and the reduced size of pOPS0359 (5.5kb) compared to pLMB509 (6.8kb) perhaps leading to an increased plasmid copy number.

### Gene expression using pOGG026

To demonstrate the flexibility of BEVA we constructed a relatively low copy number, stable plasmid pOGG026, suitable for experiments in environments where there is no antibiotic selection present.

To demonstrate the stability of pOGG026, and therefore its use for experiments carried out in many different environments, it was used to construct plasmids pOPS0253 (Fig. 2B), pOPS0254 (Fig. 2C) and pOPS0379 by Golden Gate cloning. Using a level 1 one-pot reaction (BsaI and ligation), pOPS0253 was constructed from the following (listed in Table 4): vector; pOGG026, PU module; Rlv3841pNifH from pOGG082, SC module; *gusA* from pOGG083; T module; rrnBT1 terminator from pOGG003. Plasmids pOPS0254 and pOPS0379 are identical to pOPS0253, except that the SC modules are *celB* from pOGG050 and *sfGFP* from pOGG037, rather than *gusA*. Positive controls were constructed with the constitutive neomycin cassette promoter (pNptII) from pOGG001 (PU) instead of the Rlv3841pNifH from pOGG082 resulting in the plasmids pOSP0750, pOPS0314 and pOPS0377.

Plasmids pOPS0253, pOPS0254 and pOPS0379 were then conjugated into *R. leguminosarum* Rlv3841 and used to inoculate peas. Plasmid pOPS0254 was also conjugated into R. *tropici* CIAT 899 (22) and used to inoculate bean plants.

The expression of the chromogenic marker genes in plasmids pOPS0253 and pOPS0254 can be followed by staining either pink (*gusA*) or blue (*celB*) when the appropriate substrate is supplied (20). Stained pea nodules are shown in Fig. 2E-F and stained bean nodules are shown in Fig. 2G. Even when all mature nodules showed correct staining, to further verify plasmid stability, at least three nodules were crushed from each plant, and eight independent colonies from those obtained were screened for antibiotic resistance. All colonies showed neomycin resistance, consistent with stable maintenance of plasmids derived from vector pOGG026. The inducible expression of *sfGFP* can be observed when nodules reach mature stage and just in the nitrogen fixing zone (Fig. 2H).

We have developed a plasmid-based system that can be utilized for assessing competitiveness between rhizobia which is similar to the chromosomally-encoded strategy described by Sánchez-Cañizares & Palacios (21). Using pOPS0379 we now have a non-invasive symbiosis-induced screening system which can be used to recover live rhizobia from nodules for further analysis.

### Golden Gate Level 2 cloning vectors

In the MoClo system, multigene constructs are constructed by assembling transcriptional units by Golden Gate level 1 cloning reaction into level 1 shuttle vectors that dictate the final position in a level 2 construct. Transcriptional units are then combined using a Golden Gate level 2 cloning reaction with BpiI (4). To demonstrate the flexibility of the assembly system we have designed and constructed a level 2 destination vector, pOGG216 (Fig. 4A). Vector pOGG2016 was made by a one-pot reaction for vector assembly (Esp3I and ligase) with the following modules (listed in Table 3): position 1; level 2 cloning site from pOGG006, position 2; tetracycline-resistance module from pOGG042, position 3; pBBR1 origin from pOGG011 and an appropriate Endlinker (ELT3) from pOGG013.

**FIG 4.**
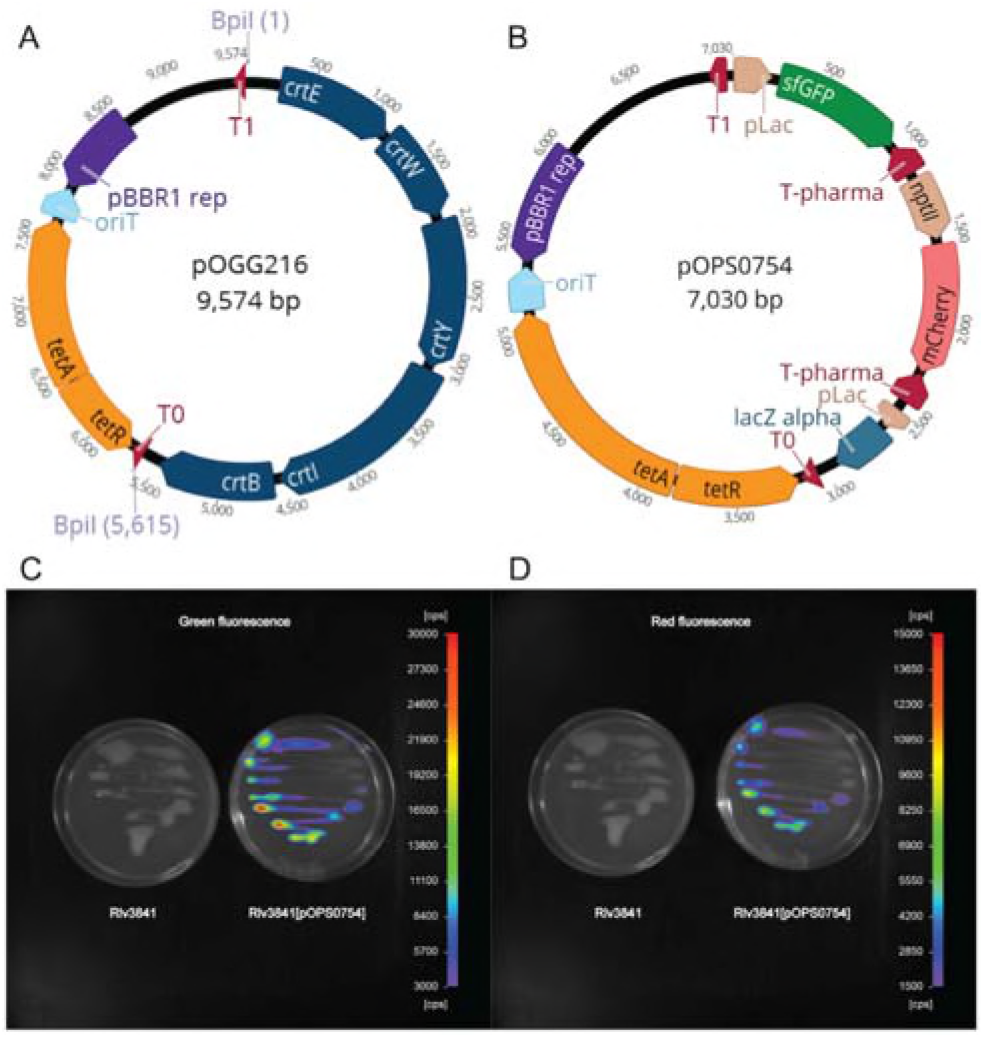
(A) Schematic map of pOGG216 level 2 broad-host range, medium copy number vector constructed with BEVA for multi-gene expression. (B) Schematic map of pOPS0754 dual reporter plasmid which encodes IPTG-inducible sfGFP cloned in Forward Position 1 and constitutively mCherry cloned in Forward Position 2. (C-D) Fluorescence of Rlv3841 and Rlv3841[pOPS0754] grown on media containing 0.5 mM IPTG. (C) Green fluorescence from IPTG-inducible sfGFP (scale, 3,000-30,000 cps). (D) Red fluorescence from constitutively expressed mCherry (scale, 1,500-15,000 cps).

### Gene expression using vector pOGG216

To validate the performance of vector pOGG216, a level 2 cloning reaction was performed (one-pot reaction with BpiI and ligase) to construct pOPS0754 (Fig. 4B), a dual reporter plasmid, using the following parts: level 2 vector pOGG216, forward position 1 from pOGG202, forward position 2 from pOGG203 and level 2 Endlinker ELB-2 from pOGG056. Plasmid pOGG202 (clone of a level 1 unit which encodes IPTG-inducible sfGFP) was constructed by combining the following in a one-pot reaction (BsaI and ligase): forward position 1 shuttle vector pL1V-F1 pOGG021, PU module pLac from pOGG031, SC module sfGFP from pOGG037 together with T module rrnBT1 from pOGG003.

Plasmid pOGG203 (clone of a level 1 unit which encodes a constitutively-expressed mCherry) was assembled by combining the following in a one-pot reaction (BsaI and ligase): forward position 2 shuttle vector pL1V-F2 pOGG054, PU module pNeo from pOGG001, SC module mCherry from EC15071 and T module rrnBT1 from pOGG003.

Assembly in *E. coli* of the final dual reporter plasmid pOPS0754 was very efficient (>90% correct constructs, same as in Weber *et al.* 2011 (4)). Plasmid pOPS0754 was conjugated into Rlv3841 and green and red fluorescence from this dual reporter plasmid is shown in Fig. 4.C-D.

In conclusion, the BEVA system we describe has proved to be robust and flexible for creating new bacterial vectors. The modular vectors we constructed with this system behaved consistently well in the Gram-negative bacteria tested (*E. coli* and *R. leguminosarum*), and demonstrated stability in the environments tested (rhizosphere and nitrogen-fixing root nodules formed by symbiotic bacteria) when engineered to contain the stability accessory module (*par* genes). The BEVA vectors for use in Golden Gate cloning (level 1: pOGG005, pOGG024 and pOGG026, and level 2: pOGG216) have been made available through Addgene (Table 2). While these vectors are useful for cloning and gene expression in *R. leguminosarum*, the major advantage of the modular system described here is its flexibility. The ability to rapidly assemble BEVA vectors with unique features, specifically suited to a specific purpose, e.g. alternative antibiotic resistance modules to avoid conflict with other markers, or to alter the origin of replication to avoid plasmid incompatibility has enormous applications for those working in many different areas. Rapid assembly of new BEVA vectors from such a bank of parts in this way offers a quick and economical option for construction of new vectors, not only for bacteria, as the system could be expanded to include parts for plant transformation or replication and maintenance in fungi.

## MATERIALS AND METHODS

### Bacterial media and growth conditions

Bacterial strains and plasmids used in this study are listed in Table 2. *E. coli* strains were grown in liquid or on solid Luria–Bertani (LB) medium (23) at 37°C, supplemented with antibiotics at the following concentrations: tetracycline (10 μg ml^−1^), gentamicin (10 μg ml^−1^), kanomycin (20 μg ml^−1^) or spectinomycin (100 μg ml^−1^). Rhizobial strains were grown on tryptone yeast (TY) agar or broth (24) or universal minimal salts (UMS)(25) at 28°C. Antibiotics were added to TY when necessary at the following concentrations: streptomycin (500 μg ml^−1^), tetracycline (5 μg ml^−1^), gentamicin (20 μg ml^−1^), rifampicin (50 μg ml^−1^), and neomycin (40 μg ml^−1^). Plasmids were transferred into wild-type (Rlv3841 and R. *tropici*
CIAT899 backgrounds by triparental mating according to Figurski and Helinski (1979) (26)

### Golden Gate Vector Assembly, level 1 and level 2 cloning

Vector assembly was performed in a thermocycler in 15 uL total volume, containing 40 fmols of each cloned module (position or ELT) (which equates to approx. 1 uL of 4 Kb plasmid at 100 ng/uL), 1.5 uL Bovine Serum Albumin (1 mg/mL), 1 uL Esp3I FD (Thermo Scientific), 1 uL concentrated T4 DNA Ligase (New England Biolabs), 1.5 uL T4 DNA Ligase buffer (New England Biolabs) with water to 15 uL. Tubes were prepared on ice before 25 cycles: 3 min at 37°C then 4 min at 16°C, followed by 5 min at 50°C and 5 min at 80°C. Samples were cooled on ice and 1 uL was transformed into 10 uL of *E. coli* competent cells (Bioline Gold) by heat shock for 1 min at 42°C (27). and plated onto LB with appropriate antibiotics (for pOGG024 selection was for gentamicin-resistance, for pOGG026 for kanamycin resistance and for pOGG216 for tetracycline resistance) and X-gal at 40 ug/mL. Several colonies showing the correct colour, i.e. blue for pOGG024 and pOGG026 (level 1) vector assembly (colour is conferred by the presence of *lacZa* in the cloning site) and orange for pOGG216 (level 2) (conferred by a canthaxanthin biosynthesis cluster in the cloning site) (4), were screened by plasmid restriction digest and sequencing to verify correct constructs. Correct plasmid assembly was observed at very high frequencies and comparable to those described for Golden Gate cloning reactions (4).

Level 1 cloning was performed as described above, except BsaI FD (Thermo Scientific) replaced Esp3I. For level 2 cloning, BpiI (Thermo Scientific) was used instead of Esp3I. Transformation was performed as described above and colonies showing the correct colour based on the replacement of modules in the vector cloning sites (blue to white for level 1 and orange to blue or white for level 2) were screened by plasmid restriction digest and sequencing to verify correct constructs.

### Plant growth

*Pisum sativum* (pea) and *Phaseolus vulgaris* (bean) seeds were surface sterilized by immersion in 95% ethanol for 0.5 min, followed by washing with sterile water. Seeds were then immersed in 2% Sodium hypochlorite for 5 min. After washing the seeds five times with sterile water, they were placed onto 1% agar (DWA) plates in the dark at room temperature for 4–5 days to allow germination. Pea seedlings were placed in sterile 1-litre pots with medium vermiculite and supplied with 400 ml nitrogen-free rooting solution (28). In the case of beans, seedlings were placed in sterile 2-litre pots with fine vermiculite and supplied with 800 ml nitrogen-free rooting solution, (29) Inoculation was at approx. 10^7^ cfu of rhizobia per pot. Plants without inoculation were grown as a control. Plants were grown (random positioning of pots) in a controlled growth chamber at 21°C, 16-h/8-h day/night cycle.

### Enzyme activity in nodules

Pea plants were harvested 21 dpi and bean plants 42dpi. Roots were stained with Magenta-glcA (5-bromo-6-chloro-3-indolyl-β-D-glucuronide acid, 200 μg/ml) for plants nodulated with strain Rlv3841[pOPS0253] containing the β-glucuronidase gene (*gusA*). For plants nodulated with rhizobial strains Rlv3841[pOPS0254] and CIAT 899[pOPS0254] containing β-glucosidase gene (*celB*), roots were stained with X-gal (5-bromo-4-chloro-3-indolyl-β-D-galactopyranoside, 250 μg/ml) after thermal treatment at 70°C for 30 minutes to destroy endogenous β-galactosidases (20).

### Measurement of bacterial growth and GFP assays

Bacterial growth OD_595_ and GFP fluorescence (excitation 485 nm and emission 520 nm) were performed with a FLUOstar OMEGA (Lite). Rhizobia were grown on TY agar slopes with appropriate antibiotics for three days, washed with UMS and resuspended in UMS with 30 mM pyruvate and 10 mM NH_4_Cl (as carbon and nitrogen sources) at an OD_595_ of 0.1 at the start of the assay. Cells were incubated at 28°C with orbital shaking at 500 RPM for 48 hours with measurement of optical density and fluorescence every 30 min.

## Microscopy

Photographs were taken using a dissecting microscope (Leica M165 FC) with a Leica DFC310 FX digital camera accompanying software (LAS v4.5). Visible light filter and a GFP filter were used.

## Image Acquisition for green and red fluorescence expression

NightOWL camera (Berthold Technologies). Each CCD image consisted of an array 1,024 by 1,024 pixels, and after acquisition, images were postprocessed for cosmic suppression and background correction. Fluorescence CCD images were acquired with a Ring-light epi illumination accessory exposed for one second and special filters have to be used. Filter used for GFP quantification at excitation wavelength 475/20 and emission wavelength 520/10 nm. Filter used for mCherry quantification at excitation wavelength 550/10 and emission wavelength 620/10 nm. Images were analysed with the imaging software IndiGO (Berthold Technologies).

## ACKNOWLEDGEMENTS

We thank Chris Rogers from Giles’s lab. Kyle Grant and Beatriz Jorrín. Carmen Sánchez-Cañizares for training MMS in the use of *gusA* and *celB* marker genes and Alison East for her critical revision of the manuscript.

This work was supported by the Biotechnology and Biological Sciences Research Council [grant number BB/L011484/1] and C0NACYT-I2T2 Nuevo León (grant code 266954 / 399852) awarded to MMS.

## REFERENCES

1. Church GM, Elowitz MB, Smolke CD, Voigt CA, Weiss R. 2014. Realizing the potential of synthetic biology. Nat Rev Mol Cell Biol 15:289–294.

2. Kosuri S, Church GM. 2014. Large-scale de novo DNA synthesis: technologies and applications. Nat Methods 11:499–507.

3. Smanski MJ, Bhatia S, Zhao D, Park Y, Woodruff LBA, Giannoukos G, Ciulla D, Busby M, Calderon J, Nicol R. 2014. Functional optimization of gene clusters by combinatorial design and assembly. Nat Biotechnol 32:1241–1249.

4. Weber E, Engler C, Gruetzner R, Werner S, Marillonnet S. 2011. A modular cloning system for standardized assembly of multigene constructs. PLoS 0ne 6:e16765.

5. Engler C, Gruetzner R, Kandzia R, Marillonnet S, Duguay A. 2009. Golden gate shuffling: a one-pot DNA shuffling method based on type IIs restriction enzymes. PLoS One 4:e5553.

6. Moore SJ, Lai H-E, Kelwick RJR, Chee SM, Bell DJ, Polizzi KM, Freemont PS. 2016. EcoFlex: A Multifunctional MoClo Kit for *E. coli* Synthetic Biology. ACS Synth Biol 5:1059–1069.

7. Terfrüchte M, Joehnk B, Fajardo-Somera R, Braus GH, Riquelme M, Schipper K, Feldbrügge M. 2014. Establishing a versatile Golden Gate cloning system for genetic engineering in fungi. Fungal Genet Biol 62:1–10.

8. Engler C, Youles M, Gruetzner R, Ehnert T-MM, Werner S, Jones JDG, Patron NJ, Marillonnet S. 2014. A Golden Gate modular cloning toolbox for plants. ACS Synth Biol 3:839–843.

9. Agmon N, Mitchell LA, Cai Y, Ikushima S, Chuang J, Zheng A, Choi W-J, Martin JA, Caravelli K, Stracquadanio G, Boeke JD. 2015. Yeast Golden Gate (yGG) for the Efficient Assembly of *S. cerevisiae* Transcription Units. ACS Synth Biol 4:853–859.

10. Martínez-Garćía E, Aparicio T, Goñi-Moreno A, Fraile S, De Lorenzo V. 2015. SEVA 2.0: An update of the Standard European Vector Architecture for de-/re-construction of bacterial functionalities. Nucleic Acids Res 43:D1183–D1189.

11. Silva-Rocha R, Martínez-García E, Calles B, Chavarriá M, Arce-Rodríguez A, De Las Heras A, Páez-Espino AD, Durante-Rodríguez G, Kim J, Nikel PI, Platero R, De Lorenzo V. 2013. The Standard European Vector Architecture (SEVA): A coherent platform for the analysis and deployment of complex prokaryotic phenotypes. Nucleic Acids Res 41:D666–D675.

12. Lutz R, Bujard H. 1997. Independent and tight regulation of transcriptional units in Escherichia coli via the LacR/O, the TetR/O and AraC/I1-I2 regulatory elements. Nucleic Acids Res 25:1203–1210.

13. Prell J, Boesten B, Poole PS, Priefer UB. 2002. The *Rhizobium leguminosarum* bv. viciae VF39 γ-aminobutyrate (GABA) aminotransferase gene *(gabT)* is induced by GABA and highly expressed in bacteroids. Microbiology 148:615–623.

14. Frederix M, Edwards A, Swiderska A, Stanger A, Karunakaran R, Williams A, Abbruscato P, Sanchez-Contreras M, Poole PS, Downie JA. 2014. Mutation of *praR* in *Rhizobium leguminosarum* enhances root biofilms, improving nodulation competitiveness by increased expression of attachment proteins. Mol Microbiol 93:464–478.

15. Berrow NS, Alderton D, Owens RJ. 2009. The precise engineering of expression vectors using high-throughput In-Fusion^TM^ PCR cloning. High Throughput Protein Expr Purif Methods Protoc 75–90.

16. Keen NT, Tamaki S, Kobayashi D, Trollinger D. 1988. Improved broad-host-range plasmids for DNA cloning in Gram-negative bacteria. Gene 70:191–197.

17. Tett AJ, Rudder SJ, Bourdès A, Karunakaran R, Poole PS. 2012. Regulatable vectors for environmental gene expression in *Alphaproteobacteria*. Appl Environ Microbiol 78:7137–7140.

18. Johnston AWB, Beringer JE. 1975. Identification of the rhizobium strains in pea root nodules using genetic markers. J Gen Microbiol 87:343–50.

19. Pédelacq J-D, Cabantous S, Tran T, Terwilliger TC, Waldo GS. 2006. Engineering and characterization of a superfolder green fluorescent protein. Nat Biotechnol 24:79.

20. Sessitsch A, Wilson K, Akkermans A, de Vos W. 1996. Simultaneous detection of different *Rhizobium* strains marked with either the *Escherichia coli gusA* gene or the Pyrococcus furiosus *celB* gene. Appl Environ Microbiol 62:4191–4194.

21. Sánchez-Cañizares C, Palacios J. 2013. Construction of a marker system for the evaluation of competitiveness for legume nodulation in *Rhizobium*strains. J Microbiol Methods 92:246–249.

22. Graham PH, Viteri SE, Mackie F, Vargas AT, Palacios A. 1982. Variation in acid soil tolerance among strains of *Rhizobium phaseoli*. F Crop Res 5:121–128.

23. Sambrook J, Fritsch EF, Maniatis T. 1989. Molecular cloning: a laboratory manual. Cold Spring Harbor.

24. Beringer JE. 1974. R factor transfer in *Rhizobium leguminosarum*. Microbiology 84:188–198.

25. Poole PS, Schofield N, Reid CJ, Drew EM, Walshaw DL. 1994. Identification of chromosomal genes located downstream of *dctD* that affect the requirement for calcium and the lipopolysaccharide layer of *Rhizobium leguminosarum*. Microbiology 140:2797–809.

26. Figurski DH, Helinski DR. 1979. Replication of an origin-containing derivative of plasmid RK2 dependent on a plasmid function provided in trans. Proc Natl Acad Sci 76:1648–1652.

27. Engler C, Kandzia R, Marillonnet S. 2008. A one pot, one step, precision cloning method with high throughput capability. PLoS One 3:e3647.

28. Poole PS, Schofiel NA, Reid CJ, Drew EM, Walshaw DL. 1994. Identification of chromosomal genes located downstream of *dctD* that affect the requirement for calcium and the lipopolysaccharide layer of *Rhizobium leguminosarum*. Microbiology 140:2797–2809.

29. Broughton WJ, Dilworth MJ. 1971. Control of leghaemoglobin synthesis in snake beans. Biochem J 125:1075–80.

